# Understanding the evolution and clade diversity of *Mayorella* (Amoebozoa): Description of Novel Species from freshwater near Cape coast, Ghana

**DOI:** 10.64898/2026.01.15.699748

**Authors:** Yonas I. Tekle, Saron Ghebezadik, Christon Jairus M. Racoma, Angelicia A. Greer, Kwaku Oti Acheampong, Kwaku Brako Dakwa

**Affiliations:** Spelman College, 350 Spelman Lane Southwest, Atlanta, Georgia 30314, U.S.A.; Department of Conservation Biology and Entomology, University of Cape Coast, Ghana

**Keywords:** *Mayorella*, Amoebozoa, Dermamoebida, freshwater amoebae, morphological plasticity, SSU rDNA phylogeny, new species, protist biodiversity, West Africa

## Abstract

*Mayorella* species are globally distributed amoeboid protists, yet their diversity remains poorly resolved due to pronounced morphological plasticity and limited molecular data. Freshwater ecosystems of West Africa are particularly underexplored, with virtually no taxonomic or molecular studies addressing *Mayorella* and most microbial eukaryotes diversity in the region. Here, we describe a new species of *Mayorella* isolated from Budamu Stream, Ghana, using an integrative morphological and molecular approach. *Mayorella budamensis* sp. nov. exhibits marked morphological plasticity in size, shape, locomotion, and floating forms. Phylogenetic analyses based on SSU rDNA place the new species within a well-supported *Mayorella* clade, clustering with geographically diverse but largely unnamed and uncultured lineages. Comparative sequence analyses confirm that *M. budamensis* sp. nov. is genetically distinct from all sequenced congeners recognized to date. In addition, a second *Mayorella* isolate recovered from the same locality forms a genetically distinct lineage associated primarily with environmental sequences, indicating a novel evolutionary group. Based on available SSU rDNA data, we recognize up to seven major clades within *Mayorella*, representing distinct evolutionary lineages with the potential to include additional undescribed species. Together, these findings reveal substantial cryptic diversity and expand the known distribution of *Mayorella* in understudied African freshwater ecosystems.

## INTRODUCTION

Amoebae of the genus *Mayorella* Schaeffer, 1926 are widely distributed across freshwater, marine, and soil ecosystems, yet their true diversity remains substantially underestimated (Schaeffer 1926, Page 1983a, Page 1983b, Bovee 1961, Cann 1981). Members of the genus play important ecological roles as polyphagous grazers of bacteria and voracious predators of other microbial eukaryotes, contributing to microbial food-web dynamics and nutrient cycling (Laybourn-Parry et al. 1987, Geisen et al. 2016). Despite their ecological significance and frequent occurrence, *Mayorella* has long been regarded as a taxonomically unstable genus (Page 1972, Page 1981, Page 1987, Glotova et al. 2018, Smirnov et al. 2011).

Historically, taxonomic studies of *Mayorella* relied almost exclusively on light-microscopy observations of locomotive and floating forms, often accompanied by schematic illustrations (Penard 1902, Schaeffer 1926, Bovee 1961). Subsequent work incorporated additional characters, including ultrastructural observations (Cann 1981, Page 1983a, Goodkov and Buryakov 1988) and limited molecular data (Dyková et al. 2008, Glotova et al. 2018, Patsyuk 2023, Lei et al. 2023, Yamagishi et al. 2025). However, species delimitation has still remained problematic due to the scarcity of stable morphological traits, pronounced morphological plasticity, and the difficulty of maintaining most strains in long-term culture (Glotova et al. 2018). Some diagnostic features, such as the fine structure of the cell coat, require electron microscopy and are available for only a small fraction of described taxa (e.g., (Page 1983a). As a result, although more than twenty nominal *Mayorella* species have been reported, many remain insufficiently documented, lack type material or ultrastructural characterization, and cannot be confidently recognized or re-isolated (Page 1983a, Page 1972, Bovee 1951, Glotova et al. 2018).

The application of molecular phylogenetics has begun to clarify this taxonomic uncertainty. Prior to 2018, a couple of small subunit ribosomal DNA (SSU rDNA; 18S) sequences were available for *Mayorella*, severely limiting phylogenetic inference (Dyková et al. 2008, Fahrni et al. 2003). Expanded SSU rDNA sampling by Glotova et al. (2018) revealed that *Mayorella* comprises multiple deeply diverging clades and harbors far greater genetic diversity than previously recognized. Many of these clades consist entirely of unnamed strains or environmental sequences, indicating that a substantial proportion of *Mayorella* diversity remains undescribed and that morphological convergence and plasticity obscure evolutionary relationships among lineages.

Current knowledge surrounding the diversity of Amoebozoa and other microbial eukaryotes is mainly influenced by pronounced geographic sampling biases. Despite global efforts to systematize every known diversity, there are still a significant portion of continents that are yet to be explored (Kudryavtsev et al. 2021). In West Africa, freshwater ecosystems remain largely untapped with respect to amoebae. Our recent surveys in southern Ghana employing culture-enriched metabarcoding and targeted isolation approaches have revealed unexpectedly high diversity of free-living amoebae across multiple amoebozoan lineages, including both pathogenic and non-pathogenic taxa (Tekle et al. 2025). During these surveys, samples collected from Budamu Stream in Cape Coast, Ghana yielded diverse mayorellid amoebae exhibiting pronounced morphological variability in locomotive form, cell size, and subpseudopodial dynamics. In this study, we describe a new species of *Mayorella* isolated from Budamu Stream using an integrative approach combining live-cell light microscopy and SSU rDNA phylogenetic analyses. We also report and phylogenetically characterize an additional Ghanaian *Mayorella* lineage known only from the same environmental sequence data. In addition, we reconstruct the most comprehensive SSU rDNA phylogeny of *Mayorella* to date using all publicly available sequences to assess evolutionary relationships and lineage diversity within the genus. Together, these findings expand the known geographic representation and genetic diversity of *Mayorella* and provide morphological and molecular data that elucidates taxonomic uncertainty and cryptic diversity in this morphologically plastic genus.

## MATERIALS AND METHODS

### Isolation, Culturing and Light Microscopy observation

Samples used for the isolation of amoebae investigated in this study were collected from Budamu Stream (5°16′27.348″ N, 1°18′32.238″ W), near Cape Coast in the Twifo Hemang Lower Denkyira District, Ghana. These samples were originally collected as part of a broader metabarcoding survey of freshwater ecosystems in the region (Tekle et al. 2025a).

Debris settled from water samples was used to initiate enrichment cultures through successive culturing and isolation steps. Small portions of the debris were transferred into plastic Petri dishes containing bottled spring water (Deer Park; Nestlé Corp., Glendale, CA) and incubated at room temperature. Cultures were supplemented with autoclaved rice grains to promote microbial growth. Amoeboid cells appearing in culture were examined using an ECLIPSE Ti2 inverted microscope (Nikon Corporation, Japan).

Isolates tentatively assigned to the genus *Mayorella* based on gross morphological features were selected for further isolation and monoclonal culturing. Because *Mayorella* species typically feed on microeukaryotic prey, establishing monoclonal cultures supported solely by bacteria proved challenging. To circumvent this limitation, individual cells were picked, cleaned, and starved prior to downstream single-cell transcriptomic analyses (see below).

Cells used for general morphological and behavioral observations were examined using the ECLIPSE Ti2 inverted microscope equipped with the NIS-Elements software package under phase-contrast and differential interference contrast (DIC) optics. The isolate described here was eventually established as a non-axenic monoclonal culture maintained on either bacteria alone or a mixed prey consortium consisting of *Naegleria clarki* plus bacteria. The majority of morphological observations and all molecular analyses presented in this study are based on this clonal line, which represents the new species described herein.

### Recovery of SSU rDNA from single-cell RNA-Seq sequencing

For single-cell transcriptomic analyses, individual amoebae were picked, thoroughly cleaned, and starved to eliminate ingested prey from food vacuoles. Cells were selected based on diagnostic morphological characteristics and photodocumented prior to further processing. Single-cell RNA sequencing followed the protocol described in Tekle et al. (2020). Between one and eight cleaned single cells were collected by mouth pipetting into sterile 0.2-mL PCR tubes and processed using the SMART-Seq v4 Ultra Low Input RNA Kit (Takara Bio USA). Sequencing libraries were prepared from 1 ng of amplified cDNA using the Nextera XT DNA Library Preparation Kit (Illumina Inc., San Diego, CA), following the manufacturer’s instructions. Library concentrations were quantified using a Qubit 3.0 Fluorometer (Life Technologies) with the DNA High Sensitivity assay. All libraries were sequenced at Azenta Life Sciences (Burlington, MA) using high-output paired-end 150-bp reads. Read processing and transcriptome assembly followed Tekle et al. (2020), resulting in the recovery of a full-length SSU rDNA sequence used for subsequent phylogenetic and divergence analyses. Oxford Nanopore sequencing was also attempted on PCR amplicons generated using both full-length and barcode SSU primers following Tekle et al. (2025a). Due to the high error rate observed in the resulting reads, Nanopore-derived SSU sequences were excluded from the final dataset.

### Phylogenetic and Sequence Divergence Analyses

Phylogenetic reconstruction and pairwise sequence divergence analyses were based on SSU rDNA (18S) sequences retrieved from public databases, including described taxa and uncultured environmental isolates showing affinity to the genus *Mayorella* based on BLAST searches and preliminary phylogenetic screening. A total of 29 ingroup taxa, including three 18S sequences (retrieved from RNA-Seq data) representing two novel isolates (YT16 and YT26) generated in this study, were analyzed together with six outgroup taxa.

SSU rDNA sequence lengths varied substantially, reflecting the inclusion of both metabarcoding-derived sequences (400–880 bp; 43%) and partial to near full-length sequences (1,520–2,934 bp; 57%). The dataset encompassed isolates from diverse habitats, including soils, freshwater environments, and shallow and deep marine systems. Geographic origins were broad and included Africa, Asia (Taiwan, China, Japan), North America, Europe, Russia, and Ukraine.

Genus-level genetic divergence was quantified using an automated Python pipeline (Tekle et al. 2025b) that processed a single FASTA file of unaligned SSU rDNA sequences. Sequences were grouped by clade number based on the phylogeny and aligned using MAFFT v7 with the L-INS-i algorithm (–localpair, –maxiterate 1000) (Katoh and Standley 2013). Pairwise proportional distances (p-distances) values were calculated exclusively among sequences within the same clade, using maximum comparable sites to account for pronounced SSU rDNA length heterogeneity. All alignments were manually inspected in AliView (Larsson 2014) to verify positional homology.

Multiple sequences alignment varying in masking percentages of ambiguous and gaps (20% (2013 bp), 50% (1346 bp), 60% (695 bp) and 80% (495 bp)) were generated using the pipeline. Phylogenetic trees using these four matrices were reconstructed in IQ-TREE (Nguyen et al. 2015, Hoang et al. 2018, Kalyaanamoorthy et al. 2017). The best-fitting evolutionary model was selected automatically with the m AUTO option. Node support was assessed using 1000 ultrafast bootstrap replicates.

## RESULT

### Morphological description of a new *Mayorella* species ‘isolate YT26’

Active cells in locomotion are elongated to irregularly triangular, with a distinctly broader anterior region and a posterior region that tapers variably (Fig.1). Measurements of actively locomoting individuals (n = 42) show that cell length ranges from ∼46–90 μm, with a mean of ∼66 μm, while breadth ranges from ∼23–58 μm, with a mean of ∼40 μm. The length-to-breadth ratio varies from ∼1–3.4 μm, with a mean of ∼1.7 μm, reflecting substantial variability associated with locomotive state. In locomotion, the cell outline is highly variable but generally elongate to irregularly triangular, with the anterior region usually broader, giving a triangular aspect typical of *Mayorella* species (Fig. 1). This outline is frequently modified during movement, and cells may transiently assume lanceolate or narrowly elongate forms, or become broadly elongate and flattened, sometimes giving the impression of a rectilinear outline with a widened anterior margin. The anterior hyaline zone is of variable length and is smooth lobed in outline (Fig. 1A– F). Despite pronounced changes in overall proportions, the cell outline remains smooth during active locomotion.

**Figure 1.**
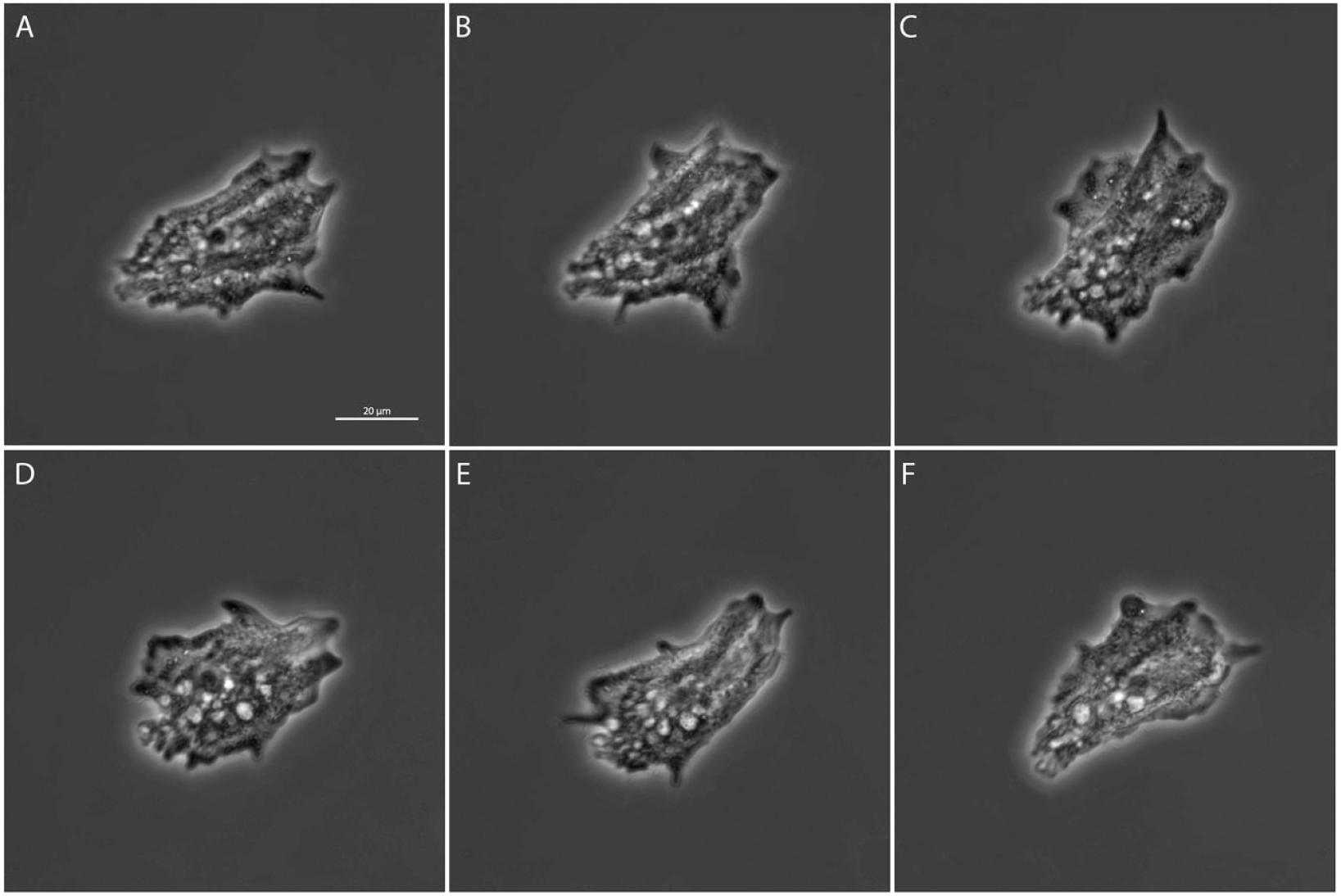
Locomotion of *Mayorella budamensis* sp. nov. in transition.

The number of subpseudopodia are variable in locomoting cells, typically ranging from 3 to 12 (Fig. 2). Forward movement may involve broad anterior adjacent subpseudopodia that extend laterally forming transient web-like hyaline sheets (Fig. 2B). In other instances, several subpseudopodia are extended simultaneously, giving the anterior region a briefly radiating appearance before one of them becomes dominant and directed movement proceeds (Fig. 2C). In rare cases, opposite ends of the cell extend simultaneously in a bidirectional manner which creates a temporary impression of a bifurcating form; then one end is subsequently retracted and unidirectional locomotion resumes (Fig. 2C). Subpseudopodia almost invariably appear first at the leading edge, projecting forward from the hyaline zone before the entire cell advances. Extended subpseudopodia originating at the anterior margin appear to trail posteriorly along the lateral margin before shortening and are resorbed near the posterior end or occur spontaneously at posterolateral. These extended subpseudopodia can be broad or tapering. The leading subpseudopodia are associated with low dorsal folds that extend to the anterior region in the granuloplasm, proportionately appearing as drawn undulations rather than discrete ridges (Fig. 2A). Although these folds are rather transient than persistent as they appear on locomoting individuals that extend their pseudopodia, whereas other individuals that glide smoothly without forming conspicuous pseudopodia, the dorsal surface appears comparatively flat. This likely indicates that they represent temporary deformations of the cell surface associated with pseudopodial activity rather than stable dorsal ridges. The posterior region lacks a clearly defined uroid in most actively moving cells but resembles closely to murolate (Fig. 1A–F). In retracting individuals, however, the posterior end may appear differentiated similar to fasciculate (Fig. 2C), though overall, it never forms a stable or sharply delimited uroidal structure. When the cell is not actively locomoting, more distinctly dactylopodial or digitiform projections may develop, particularly at the leading or posterior regions, giving the cell a more articulated appearance (data not shown).

**Figure 2.**
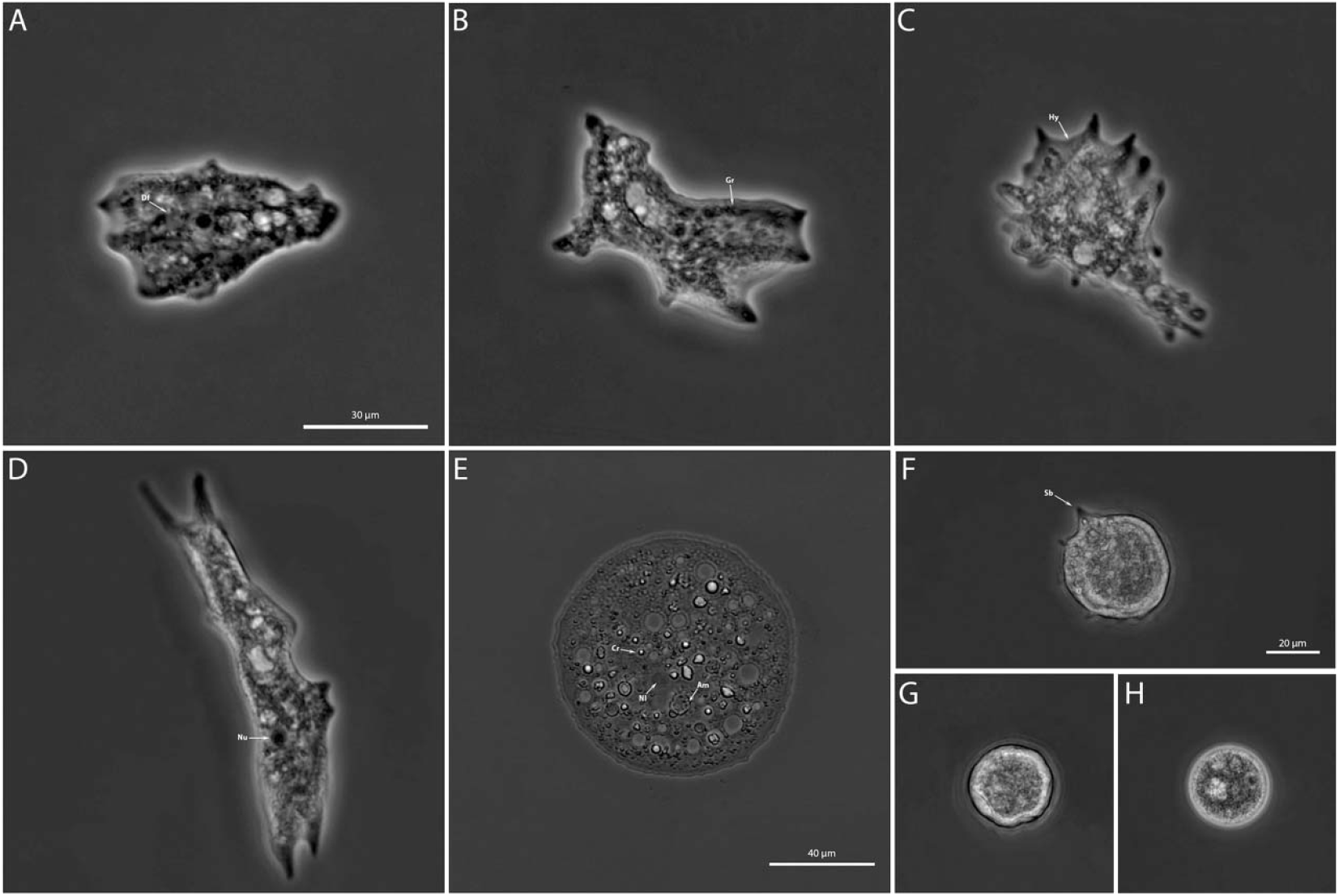
Micrographs of *Mayorella budamensis* sp. nov. A-D, displaying different morphotypes. E, floating cell mounted on a glass slide under DIC microscopy. F-G, floating forms. Df–Dorsal fold, Gr–Granuloplasm, Hy–Hyaloplasm, Nu–Nucleus, Cr–Crystal body, Nl– Nucleolus, Am–Amorphous inclusion, Sb–Subpseudopodia.

The cytoplasm is distinctly differentiated into hyaloplasm and granuloplasm. The nucleus is readily visible in the granuloplasm of locomoting cells and is typically central, though it may occasionally be displaced toward the posterior region (Supplementary Video 1). The nucleus is spherical, measuring ∼7–12 μm (n = 17) in diameter and contains a single spherical nucleolus measuring ∼5–7 μm (n = 33) in diameter (Fig. 2D, E). The granuloplasm also contains numerous vesicles and inclusions that vary widely in size and appearance. Food vacuoles are present; in some individuals, cysts of *Naegleria* sp. were observed enclosed within, indicating ingestion of eukaryotic prey (data not shown). Crystalline materials are often observed inside vesicles, while additional crystalline bodies and dark, amorphous inclusion occur freely in the granuloplasm; no consistently paired inclusions were seen. These inclusions are morphologically diverse, ranging from smooth, spherical bodies to rough, irregular ovoid forms of variable size (Fig. 2E). The contractile vacuole system consists of multiple, diffusely distributed that are variable in size and position. Individual vacuoles enlarge gradually and may fuse to the adjacent during the filling phase (Supplementary Video 1 and 2). Following discharge, the vacuolar space is rapidly replaced by surrounding granuloplasm; no distinct hyaline replacement zone was observed after emptying, and no single dominant contractile vacuole or fixed discharge site was evident.

Floating forms are present and variable in appearance. Cells frequently assume a spherical form measuring approximately ∼30–42 μm (n = 33) in diameter (Fig. 2H). Additionally, when put on cover slides, they appeared bigger in size, measuring ∼47–60 μm (n = 11). Other floating individuals appear as irregular, shriveled spheres with occasional short, broad pseudopodial extensions (Fig. 2G). In rare cases, cells were observed partially floating, with one portion of the body adhering to the substrate while the free portion bore broad, tapering pseudopodia waving (Fig. 2F). Asymmetrically radiate floating forms are rare but do occur; in these cases, long, broad tapering pseudopodia extend radially (Supplementary Video 3). Floating forms are most commonly observed in older cultures but can also be induced transiently by mechanical agitation or probing, however, cells are always transient and reverted to trophic morphology upon resumption of movement. No true cyst formation was observed.

### Molecular Data, Phylogeny and Pairwise Comparison

We present SSU rDNA sequences from two isolates designated YT26 and YT16, with GenBank accession numbers PX778840 and PX778838–PX778839, respectively. The YT26 isolate was established as a monoclonal culture and is described here morphologically as a new species of *Mayorella* (Figs. 1, 2). In contrast, the YT16 isolate was recovered only from sequence data and represents an unvouchered environmental lineage that cannot be unambiguously linked to a morphologically characterized specimen.

Phylogenetic reconstruction of the genus *Mayorella* based on SSU rDNA sequences resolved multiple distinct lineages, encompassing both formally described species and numerous unnamed or environmental taxa (Fig. 3). Across four datasets differing in the number of retained nucleotide sites, the overall topologies were largely congruent, with trees inferred from the 20% (2,013 bp) and 50% (1,346 bp) masking schemes yielding identical configurations. As the number of retained characters was progressively reduced, phylogenetic resolution declined in the 60% (695 bp) and 80% (495 bp) masked datasets; however, no strongly supported conflicts with topologies inferred from datasets containing more sites were detected (data not shown). Accordingly, the phylogeny inferred from the 20% masked dataset (2,013 bp) is presented as a representative summary (Fig. 3), and all subsequent interpretations are based on this topology.

**Figure 3.**
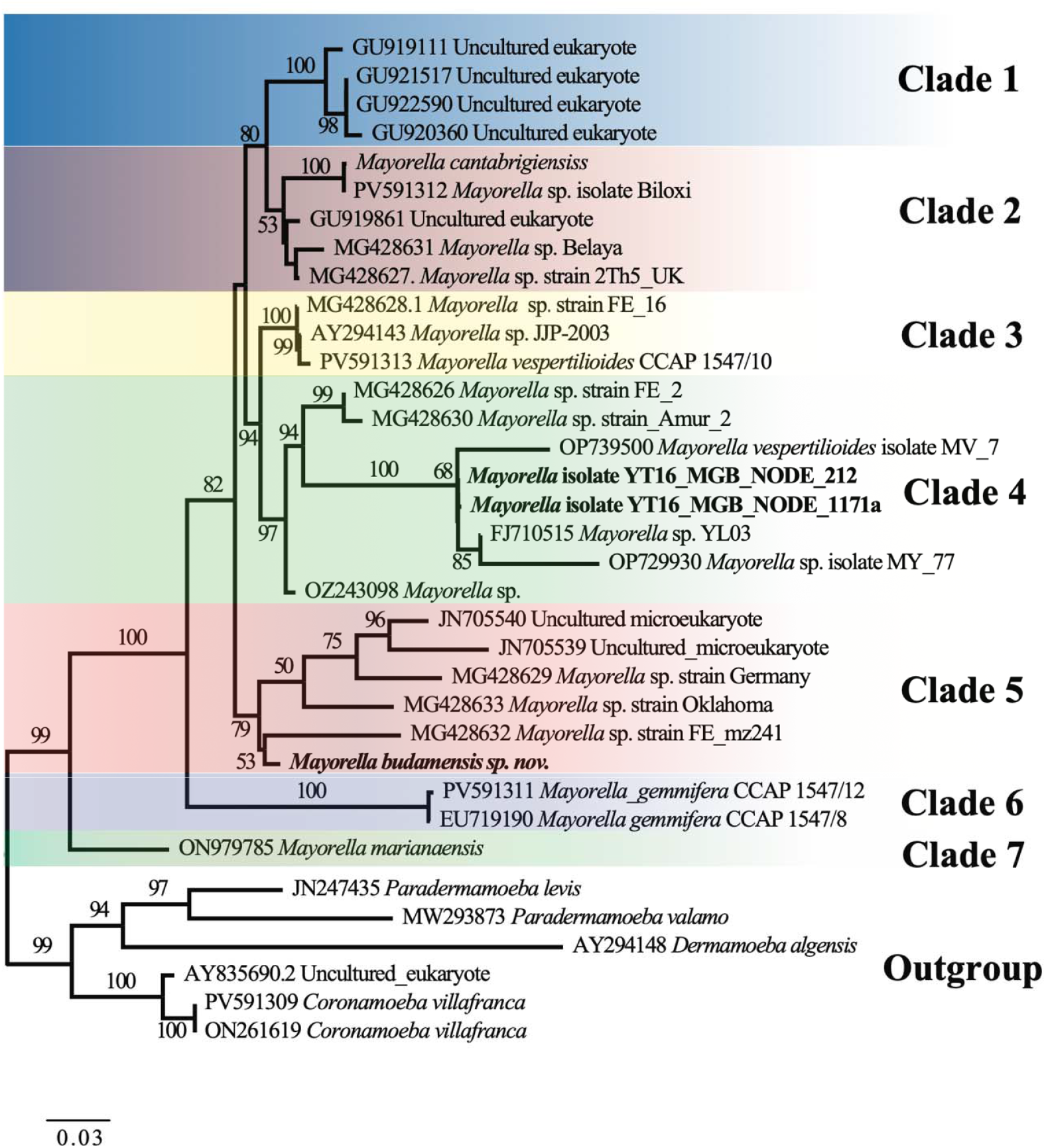
Phylogenetic placement of *Mayorella budamensis* sp. nov. within Mayorella. Maximum-likelihood tree inferred from the 20% masked SSU rDNA alignment (2,013 bp). Node support values represent ultrafast bootstrap percentages from 1,000 replicates. Scale bar indicates the expected number of substitutions per site.

The inferred phylogeny consistently recovered seven major clades, most of which received strong to moderate bootstrap support (Clades 1 and 3–7), whereas Clade 2 formed a weakly supported assemblage (Fig. 3). Pairwise p-distance values were calculated within each clade using the maximum number of comparable sites (Tables 1, S1). Across the dataset, SSU rDNA sequence length varied substantially (approximately 440–2,934 bp), reflecting the inclusion of short metabarcode-derived sequences targeting variable regions as well as partial to near full-length sequences (Table 1). This length heterogeneity complicated both overall and clade-specific distance comparisons, as many sequences overlapped only partially. Consequently, some pairwise distance estimates are based on relatively short overlapping regions and should be interpreted with caution.

**Table 1.**
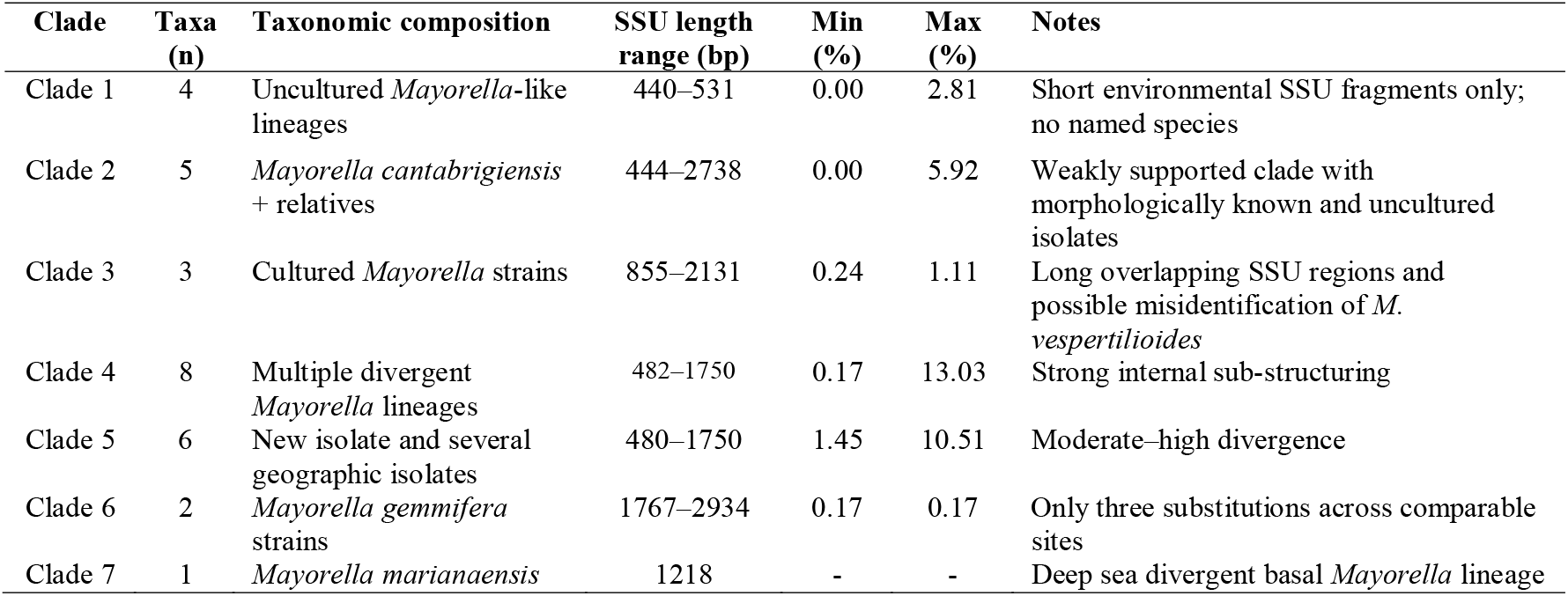
Summary of within-clade SSU rDNA divergence and sequence length heterogeneity in *Mayorella*.

### Clade 1

Clade 1 is strongly supported and consists of four uncultured eukaryotic lineages represented by short SSU rDNA sequences ranging from 440 to 531 bp (Fig. 3). Pairwise divergence within Clade 1 ranges from 0 to 2.81% (Tables 1, S1). With the exception of isolate GU919111.1 (444 bp), most observed sequence variation is attributable to indels located at the 5′ or 3′ ends, suggesting potential sequencing artefacts rather than true evolutionary divergence. Nevertheless, GU919111.1 exhibits elevated divergence (2.26–2.81%) relative to other members of the clade and may represent a distinct genetic lineage, although full-length SSU rDNA sequences will be required to evaluate this more robustly. Clade 1 is consistently recovered as the sister lineage to Clade 2 with moderate bootstrap support.

### Clade 2

Clade 2 includes five isolates comprising the named species *Mayorella cantabrigiensis* (SSU rDNA extracted from RNA-seq data), morphologically characterized isolates from Russia (MG428631.1, Belaya strain), the United Kingdom (MG428627.1, 2Th5 strain), and the United States (PV591312.1, Biloxi isolate), as well as one sequence (GU919861.2) known only from environmental data (Fig. 3). Pairwise comparisons revealed that *M. cantabrigiensis* (2,738 bp) and GU919861.2 (444 bp) are identical across their 78 bp overlapping region (Table 1); however, this apparent identity reflects the limited overlap rather than true genetic equivalence.

The Biloxi isolate (PV591312.1) and *M. cantabrigiensis* differ by only 0.063%, corresponding to one substitution and a five-base insertion across 1,581 bp of overlap (Table S1). Given the extensive shared identical regions, this low divergence likely represents intrastrain variation within the SSU rDNA marker. It should be noted, however, that the SSU rDNA sequence of *M. cantabrigiensis*, derived from RNA-seq data, lacks approximately 500 bp from the 5′ end, including the V9 hypervariable region that is commonly targeted in short-read sequencing surveys.

In contrast, the Biloxi isolate (PV591312.1) is markedly more divergent from GU919861.2 (5.923%), the Belaya isolate (MG428631.1; 4.079%), and the 2Th5 isolate (MG428627.1; 5.263%), with differences reflecting a combination of substitutions and indels across overlapping regions ranging from 444 to 881 bp (Tables 1, S1). The Belaya and 2Th5 isolates form a weakly supported sister relationship and differ by 1.05%, involving six substitutions and three indels across 576 bp of overlap. GU919861.2 differs from 2Th5 and Belaya by 1.379% and 2.069%, respectively (Table S1).

Overall, Clade 2 comprises isolates with relatively low levels of genetic divergence but includes two to three genetically distinct lineages. The Biloxi isolate and *M. cantabrigiensis* are likely conspecific or represent closely related genetic variants, given their high sequence identity across most of the SSU region (1,581 bp). However, the absence of the V9 region in the *M. cantabrigiensis* sequence complicates direct comparison, as most divergence between the Biloxi isolate and other members of the clade occurs within this region (Table S1).

### Clade 3

Clade 3 forms a fully supported monophyletic group that includes the described species *Mayorella vespertilioides* CCAP 1547/10 (Fig. 3). Pairwise divergences among MG428628.1 (855 bp; FE_16, Russia), AY294143.1 (2,131 bp; JJP-2003, Denmark), and *M. vespertilioides* (1,584 bp) are very low, ranging from 0.237 to 1.107% across comparable regions. *M. vespertilioides* is slightly more divergent than the other two isolates, differing by approximately ten substitution events (Tables 1, S1). Clade 3 is consistently recovered as the sister group to Clade 4 with strong bootstrap support.

### Clade 4

Clade 4 is the largest and most diverse lineage, consisting of eight isolates organized into three well-resolved and strongly supported subclades (Fig. 3). The first subclade comprises two unnamed Russian isolates with morphological documentation: FE_2 (MG428626.1, 482 bp) and Amur_2 (MG428630.1, 1,750 bp). These sequences are virtually identical, differing only by two indels near the end of the overlapping region (Tables 1, S1). The second subclade contains five isolates. One member, identified as *Mayorella vespertilioides* MV_7 (OP739500, 497 bp), does not cluster with *M. vespertilioides* CCAP 1547/10 and is therefore likely misidentified (Fig. 3). This isolate shows 3.823–7.930% divergence relative to other members of Clade 4 (Tables 1, S1). Two additional sequences (YT16_MGB_NODE_212 and YT16_MGB_NODE_1171a) originated from Ghanaian samples obtained during preliminary metabarcode sequencing of mixed cultures. These two sequences differ by only three nucleotides (0.174%) across 1,729 bp and are therefore considered to represent a single genetic lineage, hereafter referred to as the YT16 isolate. The YT16 isolate was not recovered in subsequent culturing efforts and is currently known only from sequence data. The YT16 isolate is 7.77–7.93% divergent from MV_7 isolate (OP739500) and likely represents a second novel *Mayorella* lineage from Ghana (Tables 1, S1).

The remaining two isolates in this subclade form a sister pair and include a Taiwanese isolate, YL03 (FJ710515.1, 516 bp), and a Ukrainian isolate, MY_77 (OP729930.1, 509 bp), which differ by approximately 20 substitutions (4.528%) (Tables 1, S1). The third subclade, forming the basal lineage of Clade 4, is represented by an environmental isolate from Japan (OZ243098.1, 688 bp). This isolate is 8.15–12.9% divergent from all other members of Clade 4 and is strongly supported as the basal lineage. Overall, pairwise divergences within Clade 4 range from 0.174 to 13.03%, indicating the presence of up to six genetically distinct species-level lineages, including the Russian MG428626.1/MG428630.1 lineages, the misidentified MV_7 isolate (OP739500), the Ghanaian YT16 lineage, the Taiwanese YL03 isolate, the Ukrainian MY_77 isolate, and the Japanese OZ243098 lineage (Fig. 3; Tables 1, S1).

### Clade 5

Clade 5 is recovered with moderate bootstrap support and includes the new YT26 isolate from Ghana (YT26_NODE_481, 1,729 bp) together with five additional isolates (Fig. 3). SSU rDNA sequence lengths within this clade range from 480 to 1,750 bp. The YT26 isolate shows low divergence from the Russian isolate FE_mz241 (MG428632.1; 1.45–1.77% across 627 bp) and the German isolate MG428629.1 (1.88–2.09% across 485 bp), based on both pipeline-generated and manually curated alignments (Tables 1, S1). These isolates are morphologically documented and display divergence values consistent with closely related strains or conspecific lineages.

In contrast, divergences between YT26_NODE_481 and the remaining members of Clade 5 are substantially higher, ranging from 4.40 to 10.51%, and exceed typical intrastrain variation, including comparisons with the morphologically characterized Oklahoma isolate MG428633 (4.399% across 684 bp) (Tables 1, S1). The remaining Clade 5 taxa include two uncultured environmental sequences from Washington State, USA (JN705540.1, 1,520 bp; JN705539.1, 1,557 bp), which exhibit 3.12–10.15% divergence across comparable regions (Tables 1, S1). Collectively, Clade 5 likely comprises five to six distinct genetic lineages, including a low-divergence group requiring additional markers or longer SSU rDNA regions for resolution (YT26_NODE_481, MG428629.1, MG428632.1) and more divergent lineages exceeding typical intrastrain thresholds (JN705540.1, JN705539.1, MG428633).

### Clades 6 and 7

Clade 6 includes two strains of *Mayorella gemmifera* (CCAP 1547/12, PV591311.1; 1,767 bp and CCAP 1547/8, EU719190.1; 2,934 bp), which differ by only three nucleotide substitutions, consistent with intraspecific variation (Fig. 3, Tables 1, S1). Clade 7 is represented by a single species originally deposited as *Dermamoebida* sp. strain 1 (ON979785.1, 1,218 bp), which forms a robustly supported basal lineage within *Mayorella* (Fig. 3). This taxon has since been formally described as *Mayorella marianaensis*, a highly divergent species isolated from deep-sea environments in the Pacific Ocean (Lei et al. 2023).

## DISCUSSION

### *Mayorella* as a ubiquitous yet poorly resolved amoeboid lineage

Species of *Mayorella* are among the most frequently encountered and easily recognized naked amoebae in freshwater, soil, and marine environments, including habitats ranging from shallow coastal zones to deep sea systems exceeding 3,000 m depth (Bovee 1951, Bovee 1961, Dyková et al. 2008, Glotova et al. 2018, Lei et al. 2023, Page 1981, Page 1983b, Page 1983a, Patsyuk 2023, Schaeffer 1926, Sawyer 1971, Bovee 1965). This widespread ecological and geographical distribution contrasts sharply with the limited taxonomic resolution of the genus, in which species level identifications have frequently been characterized by uncertainty (Glotova et al. 2018). Despite their ecological importance as active predators of bacteria, fungi, and other microbial eukaryotes, *Mayorella* remains one of the most taxonomically and phylogenetically problematic genera of naked amoebae (Page 1987, Glotova et al. 2018).

This long-standing paradox of high ecological prominence coupled with weak taxonomic resolution reflects both historical and methodological limitations that have persisted for nearly a century. Traditional taxonomy of *Mayorella* relied primarily on light microscopy observations of locomotive and floating forms, following classical treatments by Page and other early investigators (Page 1976, Page 1983a, Page 1988, Lee et al. 1985, Bovee 1951, Bovee 1970). Characters such as overall cell outline, degree of dorsoventral flattening, development of dorsal folds, number and shape of subpseudopodia, and floating morphology were widely used for species delimitation. However, experimental and comparative studies have demonstrated that these traits are highly plastic and strongly influenced by culture conditions, feeding strategies, substrate interactions, and physiological state (Glotova et al. 2018, Susan 2004). As a result, many characters historically treated as diagnostic are transient and inconsistently expressed, leading to substantial overlap among species descriptions and contributing to persistent taxonomic instability. Although ultrastructural features of the cell coat have proven to be more reliable taxonomic characters, yet such data are available for only a small fraction of described *Mayorella* species (Cann 1981, Dyková et al. 2008, Page 1983a) and their boarder taxonomic implications remain uncertain (Glotova et al. 2018).

The incorporation of molecular data has begun to address some of these limitations, but significant challenges remain. Until relatively recently, *Mayorella* was represented by only a very limited number of SSU rDNA sequences, providing insufficient phylogenetic resolution for robust species delimitation (Dyková et al. 2008, Glotova et al. 2018, Patsyuk 2023, Lei et al. 2023, Yamagishi et al. 2025). Even with expanded sampling, most available sequences derive from uncultured environmental surveys or metabarcoding studies and often target short hypervariable regions of the 18S gene (Geisen et al. 2016). As demonstrated in the present study, this heterogeneity in sequence length and genomic coverage complicates phylogenetic reconstruction and distance based comparisons, limiting the interpretive power of single marker datasets when considered in isolation. *Mayorella* therefore exemplifies a broader challenge in protist systematics, wherein neither morphology alone nor single marker molecular data are sufficient to capture true evolutionary diversity.

The present study further illustrates these limitations by demonstrating that even newly characterized *Mayorella* isolates exhibit morphological features that overlap with those reported for other described species and that vary under different culture conditions (Glotova et al. 2018). Such overlap underscores the limited discriminatory value of morphology in isolation and highlights the necessity of integrating morphological observations with molecular phylogenetic context. Conversely, molecular data lacking associated morphological documentation provide limited insight into phenotypic expression and ecological relevance.

Importantly, the *Mayorella* isolate described here represents the first formally documented and molecularly characterized species of the genus from the African continent. A previous study reported a *Mayorella* species from Uganda based on morphological and biochemical observations (Leonardi et al. 1995). Together, these observations indicate that *Mayorella* has a broad geographic distribution, occurring across diverse habitats and multiple continents, while remaining poorly characterized in large regions such as Africa. The present study helps to address this gap by providing integrative morphological and molecular data, underscoring the importance of expanded sampling in understudied regions for improving our understanding of *Mayorella* diversity, biogeography, and evolution.

### Clade diversity and cryptic species within the genus *Mayorella*

Congruent with a previous study (Glotova et al. 2018) our expanded phylogenetic reconstruction and pairwise divergence analyses reveal that *Mayorella* comprises at least seven major clades, most of which likely encompass multiple species-level lineages. Importantly, this diversity is unevenly distributed across the phylogeny and shows no consistent relationship with the availability of morphological data, underscoring the pervasive nature of cryptic diversity within the genus. The results demonstrate that both deeply divergent and closely related lineages coexist within *Mayorella*, often without clear morphological differentiation (Fig. 3).

Clade 1 illustrates the limitations of environmental sequencing for species delimitation (Fig. 3). This clade consists exclusively of uncultured lineages represented by short SSU rDNA fragments. Although overall divergence within the clade is low, one lineage exhibits elevated divergence relative to the others, suggesting the possible presence of additional species-level diversity. The absence of full-length sequences and complementary molecular markers for this clade highlights a recurring limitation of environmental surveys, in which short amplicons can obscure both true diversity and phylogenetic relationships (Pawlowski et al. 2014).

Clade 2 exemplifies a different but equally important challenge. This weakly supported assemblage includes the named species *Mayorella cantabrigiensis*, morphologically characterized isolates from Europe and North America, and uncultured environmental sequences (Tables 1, S1). Apparent sequence identity between some members reflects extremely limited overlap rather than true genetic equivalence, whereas other comparisons reveal divergences exceeding typical intrastrain variation (Tables 1, S1). These patterns indicate that Clade 2 likely harbors two to three genetically distinct lineages despite superficial morphological similarity (Glotova et al. 2018). This finding highlights the risk of equating short-region SSU rDNA identity with conspecificity and underscores the need for caution when interpreting partial sequences

In contrast, Clades 3 and 6 show relatively low levels of divergence and are consistent with single species or closely related strain complexes, as exemplified by *Mayorella vespertilioides* CCAP 1547/10 and *M. gemmifera*, respectively (Fig. 3). These clades provide useful reference points for interpreting divergence thresholds within the genus, illustrating patterns consistent with intraspecific or near-intraspecific variation in SSU rDNA.

Clades 4 and 5, however, display pronounced internal structuring and represent the clearest evidence for extensive cryptic diversity within *Mayorella*. In these clades, pairwise divergences frequently exceed values typically associated with intraspecific variation in Amoebozoa (Nassonova et al. 2010, Smirnov et al. 2007, Glotova et al. 2018). Clade 4 alone includes multiple deeply divergent lineages, several of which are known only from sequence data, including environmental and metabarcode-derived sequences from geographically distant regions. Similarly, Clade 5 contains a mixture of morphologically documented isolates, uncultured environmental lineages, and the newly described Ghanaian isolate, *Mayorella budamensis* sp. nov., with divergence patterns consistent with the presence of five to six distinct genetic lineages.

The placement of *Mayorella budamensis* sp. nov. within Clade 5 further illustrates the decoupling of morphology and evolutionary lineage within the genus. Despite pronounced morphological variability, SSU rDNA analyses consistently recover this taxon as a well-supported and genetically distinct lineage. This demonstrates that substantial phenotypic variation can occur within a single evolutionary entity and should not be interpreted as evidence for multiple species in the absence of molecular support. Conversely, morphologically similar isolates historically assigned to the same species may be distributed across multiple clades. A clear example is provided by isolates identified as *M. vespertilioides*, including MV_7, whose placement outside the clade containing the CCAP type strain strongly suggests misidentification driven by convergent morphology rather than close evolutionary relationship.

*Mayorella budamensis* sp. nov. (Ghana, Africa) shows strong morphological similarity to the European *Mayorella* strain Germany and the Russian isolate FE_mz241, including broadly triangular to elongate locomotive forms with a prominent anterior hyaline zone, variable numbers of subpseudopodia (often several extended at the leading edge), weak or absent differentiated uroidal structures, and comparable nuclear architecture (a single spherical nucleus with a central nucleolus) as well as similar floating morphologies with radiating, conical pseudopodia. This close morphological correspondence is consistent with low SSU rDNA divergence between *M. budamensis* sp. nov. and FE_mz241 (MG428632.1; 1.45–1.77% across 627 bp) and between *M. budamensis* sp. nov. and the German isolate (MG428629.1; 1.88–2.09% across 485 bp). Together, these values place the three isolates near or below commonly applied species-level divergence ranges for amoebozoans and support the interpretation that they represent a single widely distributed species or a complex of very closely related geographic variants spanning Africa, Europe, and Russia.

The presence of deeply divergent yet often uncultured lineages across multiple clades highlights the magnitude of cryptic diversity within *Mayorella*. At the same time, these results underscore the limitations of relying on SSU rDNA alone, particularly when sequence length and genomic coverage vary substantially among taxa. While SSU data provide a critical framework for recognizing major evolutionary lineages, additional molecular markers and genome-scale data will be required to fully resolve species boundaries and to integrate genetic diversity with morphological and ecological traits in this taxonomically challenging genus.

**s**

#### Taxonomic Appendix

Based on Adl et al. (2019): Amorphea: Amoebozoa: Discosea: Flabellinia Smirnov et al. 2005; Dermamoebida Cavalier-Smith 2004; *Mayorella* Schaeffer, 1926; *Mayorella budamensis* sp. nov. Tekle 2025 (Figs. 1, 2)

#### Diagnosis of *Mayorella budamensis* sp. Nov

Locomotive cells elongate to irregularly triangular with length of ∼46–90 μm, breadth of ∼23–58 μm, length-to-breadth ratio ∼1–3.4 μm. Subpseudopodia variable from broad tapering to conical or mammiliform; trailing posterior subpseudopodium consistently present during locomotion. Dorsal folds are evident on anterior ends in cells showing active pseudopodial locomotion and are not observed in flattened individuals moving by smooth gliding. Uroid weakly differentiated, occasionally approaching morulate type. Nucleus spherical, ∼7–12 μm in diameter, with single nucleolus ∼5–7 μm. Crystal and amorphous inclusions are diverse including distinctly spherical and variable ovoid shapes, often associated in vesicles or granuloplasm. Contractile vacuoles are multiple and diffuse towards the mid-posterior and may conjoin from adjacent ones. Floating form is variable but commonly spherical with instances of subpseudopodia extensions, rarely radiating. Cysts not observed. Type locality: Cape Coast, Ghana.

### Distinguishing feature of *Mayorella budamensis* sp. nov

*Mayorella budamensis* sp. nov. is characterized by pronounced morphological plasticity combined with a suite of stable cellular features that place it within a closely related lineage of freshwater *Mayorella*. Actively locomoting cells exhibit frequent transitions among elongate, triangular, and broadly elongate forms, with a smooth but prominent anterior hyaline zone, low and transient anterior dorsal folds associated with pseudopodial activity, and a posterior region that is weakly differentiated, resembled close to morulate and fasiculate but never forms a stable uroid. Subpseudopodia are variable in number and arrangement, and trailing posterior extensions may occur but are not fixed structural features. The cytoplasm contains diverse crystalline inclusions often associated with vacuoles, and the contractile vacuole system is diffuse and multi-vacuolar, lacking a distinct hyaline replacement zone after discharge. Floating forms are highly variable and include spherical, irregular, partially adherent, and, more rarely, asymmetrically radiate morphologies. These morphological traits closely match those reported for the German *Mayorella* isolate (MG428629.1) and the Russian isolate FE_mz241 (MG428632.1) and are concordant with low SSU rDNA divergence between *M. budamensis* sp. nov. and FE_mz241 (1.45–1.77% across 627 bp) and between *M. budamensis* sp. nov. and the German isolate (1.88–2.09% across 485 bp). Together, the strong morphological correspondence and minimal genetic divergence suggest that *M. budamensis* sp. nov. and these European and Russian isolates represent either a single widely distributed species or a complex of very closely related geographic variants exhibiting high cryptic diversity across Africa, Europe, and Russia, with definitive species boundaries requiring additional molecular markers and genome-scale data.

### Type locality

Freshwater sediment and detritus from Budamu Stream, near Cape Coast, Twifo Hemang Lower Denkyira District, Central Region, Ghana (5°16′27.348″ N, 1°18′32.238″ W). The stream is a lowland freshwater system characterized by slow-moving water, organic-rich sediments, and abundant microbial prey.

### Type material

#### Holotype

A monoclonal culture derived from a single isolated trophic cell collected from the type locality and established in laboratory culture. The holotype strain is preserved in cryogenic storage in the Tekle Laboratory Culture Collection, Department of Biology, Spelman College, Atlanta, Georgia, USA.

#### Hapantotype material

Photodocumented live cells representing locomotive and floating morphologies, including differential interference contrast (DIC), and phase-contrast as well as video recordings documenting locomotive behavior and morphological plasticity are archived with the corresponding author.

#### Genetic reference material

The SSU rDNA sequences derived from single-cell RNA-seq are deposited in GenBank under accession numbers PX778838-PX778840 and constitutes the molecular reference for the species.

### Etymology

The species epithet *budamensis* (Latinized adjective) refers to Budamu Stream, the freshwater system in Ghana from which the species was isolated.

#### Zoobank registration

*Pending…*

## Data Availability

The SSU rDNA sequence generated in this study is deposited in GenBank under an accession numbers PX778838-PX778840. All alignments, divergence-analysis scripts, and phylogenetic datasets used in this study are available from the corresponding author upon request. Additional materials, including raw microscopy files, will be made available through a public data repository upon request. Video documentation of cellular behavior is available through the principal investigator’s YouTube channel and upon request.

## Acknowledgments

This work was supported by the Simons Foundation Fellow Award (SFA-23-5) and the National Science Foundation Excellence in Research (EiR) Award #2401946 to Y.I.T. We thank the members of the Department of Biology at the University of Cape Coast for their assistance with fieldwork. We are also grateful to Priyal Patel for help with laboratory work.

## Supplementary Materials captions

**Table S1**. Pairwise SSU rDNA divergence within *Mayorella* clades. Pairwise SSU rDNA p-distance comparisons for each recovered *Mayorella* clade. Distances are reported as percent divergence (%) unless otherwise indicated. All calculations were performed using maximum comparable sites to accommodate SSU length heterogeneity.

**Supplementary Video 1**. Locomotion *Mayorella budamensis* sp. nov. isolate in a chambered glass slide documented under phase-contrast (10 frames per second at 40x magnification).

**Supplementary Video 2**. Floating/Resting form of *Mayorella budamensis* sp. nov. in a cover slide documented under phase-contrast (10 frames per second at 40x magnification).

**Supplementary Video 3**. Floating/Resting form (radiating) of Mayorella budamensis sp. nov. in a culture petri dish documented under an inverted microscope (40x magnification).

